# A Cooperative Model for Symmetric Ligand Binding to Protein Fibrils

**DOI:** 10.1101/2025.01.29.635590

**Authors:** Matthew S. Smith, William F. DeGrado, Michael Grabe, Brian K. Shoichet

## Abstract

A hallmark of neurodegenerative diseases like Alzheimer’s Disease (AD) and chronic traumatic encephalopathy (CTE) is the presence of toxic protein aggregates in neurons. In AD and CTE specifically, the protein tau forms insoluble fibrils that are hundreds of nanometers in length. Intriguingly, recent experimental structures suggest that tau ligands like the disaggregator EGCG and positron emission tomography (PET) tracers like GTP-1 and MK-6240 bind to tau fibrils in long stacks reflecting the symmetry of the protein across many binding sites. In these stacks, each ligand makes more contact with its symmetry mates than it does with the protein. To interpret the binding of these molecules and new ligands, we must understand the effects of the cooperativity between sites and the entropy coming from the number of sites. Here, we investigate a nearest-neighbors model of cooperativity and use statistical mechanics to derive binding isotherms for saturation and competition experiments. This model allows us to relate measured *EC*_50_ so *IC*_50_ and so values to the intrinsic binding affinity to a single site and to cooperativity across sites in ways resembling the Cheng-Prusoff Equation. Depending on the degree of cooperativity between molecular species, this model permits solutions that lack the steep binding curves expected from cooperative systems and even solutions resembling 2-site systems. We finally consider conditions for a fibril’s detection in a PET scan and practical matters of fitting this model’s parameters to data.

## Introduction

Many neurodegenerative diseases involve the accumulation of toxic protein fibrils, and clinicians can use their buildup diagnostically. For example, (PET) tracers can be used to diagnose the accumulation of fibrils from different tauopathies, like those in Alzheimer’s Disease (AD), chronic traumatic encephalopathy (CTE), progressive supranuclear palsy (PSP), and corticobasal degeneration (CBD).[1–19] These tracers have reported binding constants (*K*_*D*_ for saturation assays or *K*_1_ for competition assays) ranging from the low nanomolar to high femtomolar.

For such potent binding, both the PET tracers and their fibril binding sites are surprisingly simple. The ligands are often relatively small with few obvious binding moieties. For example, the PET tracers MK-6240, GTP-1, and flortaucipir all have several conjugated aromatic rings with one, or in the case of GTP-1, no hydrogen bond donors (**Fig. 1a**).[20] The fibril sites are also unassuming: the open “trough” of AD paired helical filament (PHF) tau has some charged and polar residues on each protein monomer but are otherwise flat, hydrophobic, and solvent-exposed (**Fig. 1b**).[21] Conversely, ligands that bind tightly to proteins typically do so in high-curvature pockets that feature pre-organized side chains that interact with multiple polar and non-polar ligand groups. Both on the fibril side and on the tracer side, such high-affinity features are lacking.

**Figure 1.**
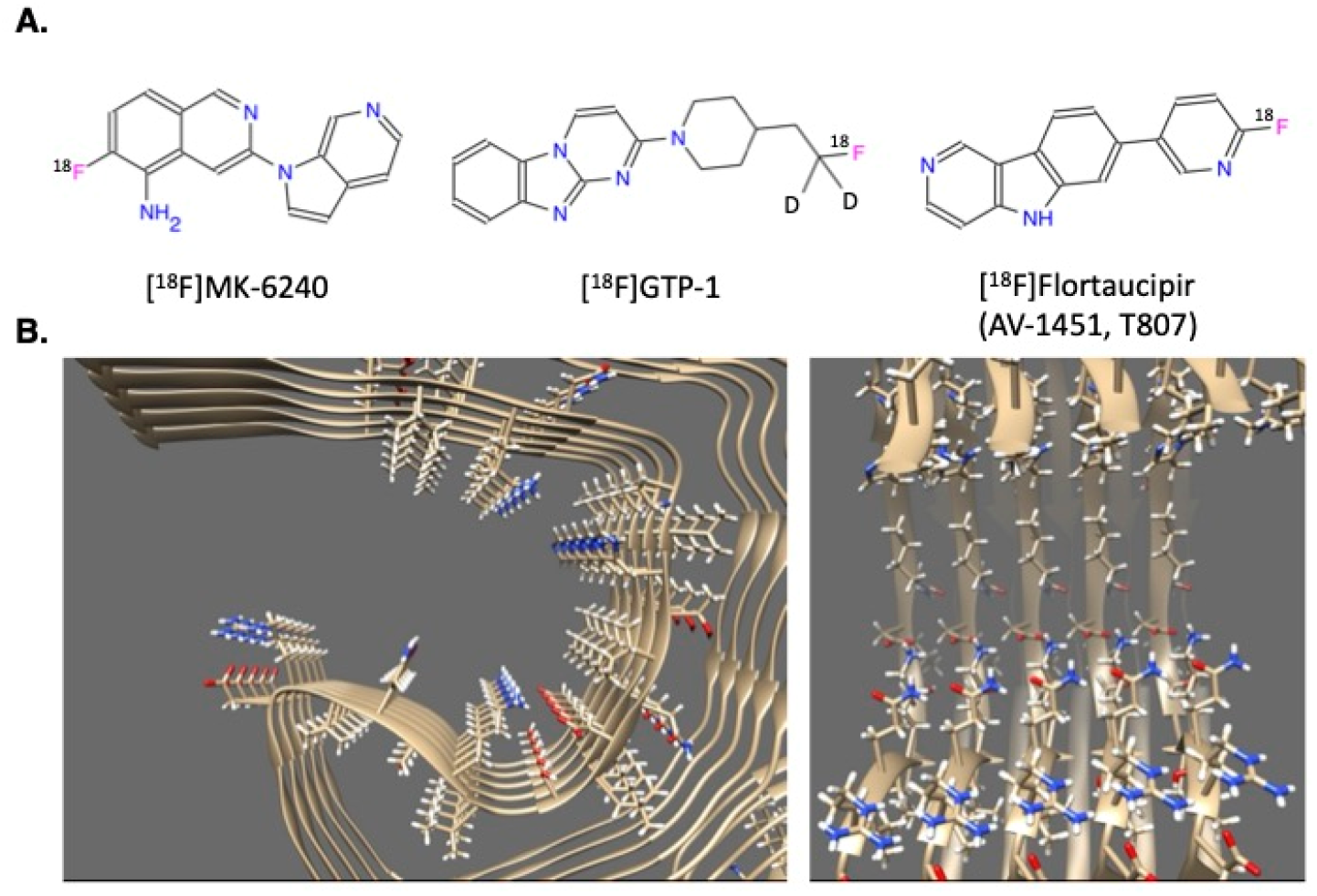
Examples of high-affinity tau ligands. (**A**.) and two views of the cryo-EM structure of an AD PHF tau protofilament (**B**.; PDB ID: 5O3L).

A potential resolution to the puzzle of fibril binding came with the determination of cryogenic electron microscopy (cryo-EM) structures of ligand-fibril complexes.[19, 22–24] In these structures, the ligands bind in long stacks with a 1:1 stoichiometry to each tau monomer and the ligand stack following the symmetry of the protein fibril. Each ligand in the stack makes few contacts with the protein and instead makes extensive contact with their ligand symmetry mates. A characteristic example is MK-6240 (right- most molecule in the top row of **Fig. 1a**) in complex with tau neurofibrillary tangles(NFTs) from AD patients (believed to bind to the AD PHF fold shown in **Fig. 1b**). In competition assays against *ex vivo* brain tissue, MK-6240 has a reported *K*_1_ of 360 pM.[10] In the structure of MK-6240 bound to AD PHF tau, however, the molecule makes only one hydrogen bond to the protein and buries more solvent-accessible surface area (SASA) in ligand-ligand interactions (243 Å^2^) than with the fibril (208 Å^2^).[23] In this complex and the complexes of GTP-1 and flortaucipir bound to their respective tau polymorphs, the ligand-ligand planar distance ranges from 3.3 Å to 3.5 Å, which is optimal for π-π interactions.[19, 22, 23] The answer to the puzzle of how such simple molecules could interact so potently with such flat sites might be that much of their affinity comes from ligand-ligand interactions.

This apparent resolution raises a second dilemma, however: from such highly interacting ligand sites, one would expect to see cooperative binding curves. This cooperativity could be positive, like oxygen binding to hemoglobin, or negative, as with tafamids binding to transthyretin.[25, 26] In either case, we would expect the effect on the curves’ steepness to be strong based on the degree of ligand-ligand interaction. But in concentration-response curves of multiple PET tracers to multiple fibrils, binding appears to follow a single-site, non-cooperative model, with a Hill Coefficient of 1.[14, 27] In some radioligand competition assays, the binding curve even appears biphasic, indicating negative cooperativity or the existence of two nonequivalent binding sites.[13, 18] While secondary binding is occasionally observed in amyloid fibrils, the density is typically too weak to be well-modeled by ligand binding, and negative cooperativity is unlikely based on the favorable ligand-ligand interactions. There is always the possibility that the cryo-EM structures, with their direct ligand-ligand interactions across every site, are artifacts of using saturating ligand concentrations, far above the concentrations used *in vivo* in PET. But this would leave us to explain where the potent binding of PET tracers comes from, our original dilemma.

Here we seek to reconcile the high-affinity binding, the apparent cooperativity in the cryo-EM structures, and the shallow concentration response curves with a new model for cooperative small molecule binding to protein fibrils. The model is based on a lattice gas framework (a modified Ising magnet), with an intrinsic affinity of the ligand to each site, cooperativity between nearest neighbors, and thousands of equivalent sites in a linear array. We use this model to make several predictions about symmetric fibril binding, like the relationship between *K*_*D*_ and *EC*_50_ in saturation assays, the relationship between *K*_*D*_ and *IC*_50_ in competition assays, and how certain forms of cooperativity in competition assays can lead to shallow or biphasic binding curves. Even in the absence of cooperativity, the model explains how the many binding sites on fibrils lowers the effective affinity needed for a tracer to be detectable in PET scan. We also suggest prospective experiments to test the model and its implications for ligand discovery.

### Description of Model

We normally think about ligands binding to proteins using the reaction

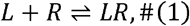

where *L* is the ligand (e.g., small molecule, peptide, protein, among others), *R* is the protein receptor (or enzyme), and *LR* is the complex. If the concentration of free ligand is [*L*] and the binding has a dissociation constant *K*_*D*_ then the Gibbs free energy change of this reaction is

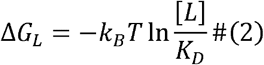

for temperature *T* and Boltzmann’s constant *K*_*B*_.[28, 29] Assuming [*L*] does not significantly change upon the binding of a small fraction of ligand molecules (i.e. no ligand depletion), then the equilibrium fraction of receptor sites bound is

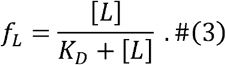

This framework assumes that every protein receptor is independent of the other ones, without any cooperativity.

Cryo-EM structures of PET tracers bound to tau show many identical binding sites in sequence. Negative-staining EM indicates that the fibrils are hundreds of nanometers long, so we conclude that there are thousands of equivalent binding sites per fibril. The simplest model of cooperativity is one of nearest neighbors, like an Ising Model of a magnet or a lattice gas.[30–33] For a given configuration of bound and unbound sites on the fibril, having 2 bound sites next to each other adds an additional energy *C*_*L*_ to the system (**Fig. 2, Table**). More negative cooperativity energies mean more favorable interactions. This cooperativity energy considers direct ligand-ligand interactions (such as Π-Π stacking, van der Waals forces, and the hydrophobic effect) as well as protein-ligand interactions, such as how the change in the ligands’ poses affect their interactions with the protein (**Fig. S1**).[34] The energy contributions are additive: if there are two molecules binding next to each other in an otherwise free stretch of the fibril, the system’s energy changes by 2Δ*G*_*L*_ + *C*_*L*_, for example.

**Figure 2.**
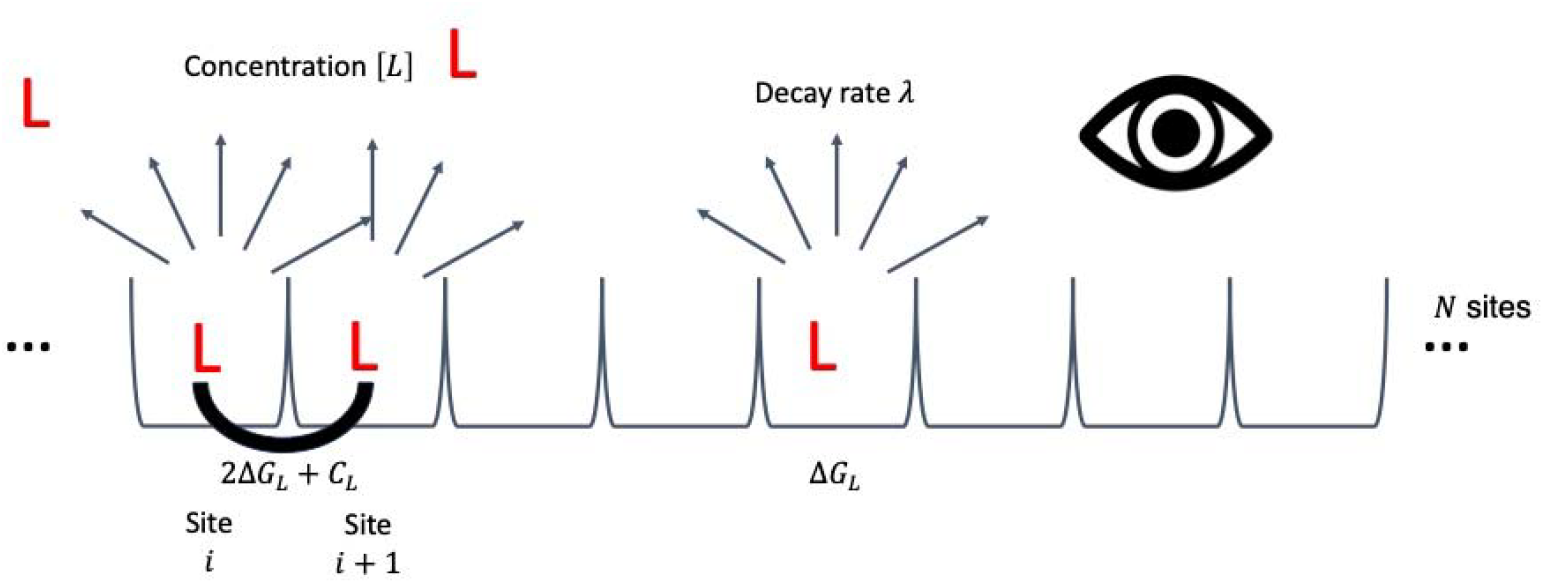
Model for cooperative ligand binding across symmetric binding sites on a protein fibril. Each protein fibril has *N* equivalent binding sites in a linear array. Each ligand molecule can bind to an individual site with dissociation constant *K*_*D*_. Assuming no ligand depletion, the free ligand concentration remains [*L*] before and after binding. For each radioligand bound, the system’s energy changes by 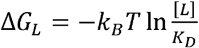 for absolute temperature *T* and Boltzmann’s constant *K*_*B*_. Every time two radioligands bind at neighboring sites, the system’s energy further changes by *C*_*L*_. The radioligands decay by beta particle or positron emission with intrinsic decay rate *λ*, which the detector can measure.

Finally, we can relate the degree of radioligand binding to the protein fibril, in its many configurations, to whether the fibril appears on a detector for beta decay (for tritiated molecules) or positron emission (for PET tracers). Each radioligand has an intrinsic decay rate *λ* (related to its half-life *t*_½_ by 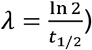. At any given time, the configuration of a fibril of length *N* has *n*_*L*_ radioligands bound and a fraction of sites bound 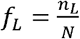. The signal (decay events per unit time) for the whole fibril is then *λn*_*L*_ =*λNf*_*L*_ We assume that ligand binding, and inhibitor binding for competition assays, happens faster than the radiation detection (or else that there are enough copies of the fibril that we can get the correct thermodynamic average between them). Then the signal on the detector (measured in counts per minute *CPM*) should be proportional to the thermodynamic average of the signal over all the configurations, with constant of proportionality *k*:

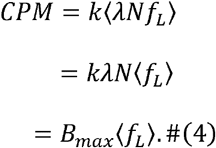

We have combined all the linear constants into *B*_*max*_, which is the maximum signal level for a given assay or PET scan setup. Based on EQ 4, we need to understand the thermodynamic average ⟨*f*_*L*_⟩ to derive a binding curve or conditions for PET detection.

In a saturation assay, each of the *N* sites on the fibril can be occupied by a radioligand or empty (we assume non-specific binding is linear with ligand concentration and its signal can be subtracted off). There are then 2^*N*^ configurations for the fibril, each one with an equilibrium probability related to its energy by Boltzmann’s relation. This system has entropy from its equivalent binding sites: many configurations have the same number of sites occupied, so we must calculate ⟨*f*_*L*_⟩ over all these configurations. Using the method of the transfer matrix (**Supplemental Methods**), we can evaluate this average in the limit of infinite sites:[35–39]

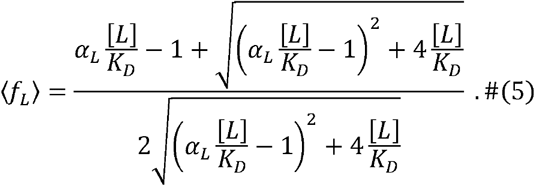

We plot this function for different cooperativity factors *α*_*L*_ on the logarithmic scale in **Fig. 3** and compare this theoretical result to Markov Chain Monte Carlo (MCMC) simulations of fibrils of finite length *N* = 1000 in **Fig. S2**.[40, 41]

**Figure 3.**
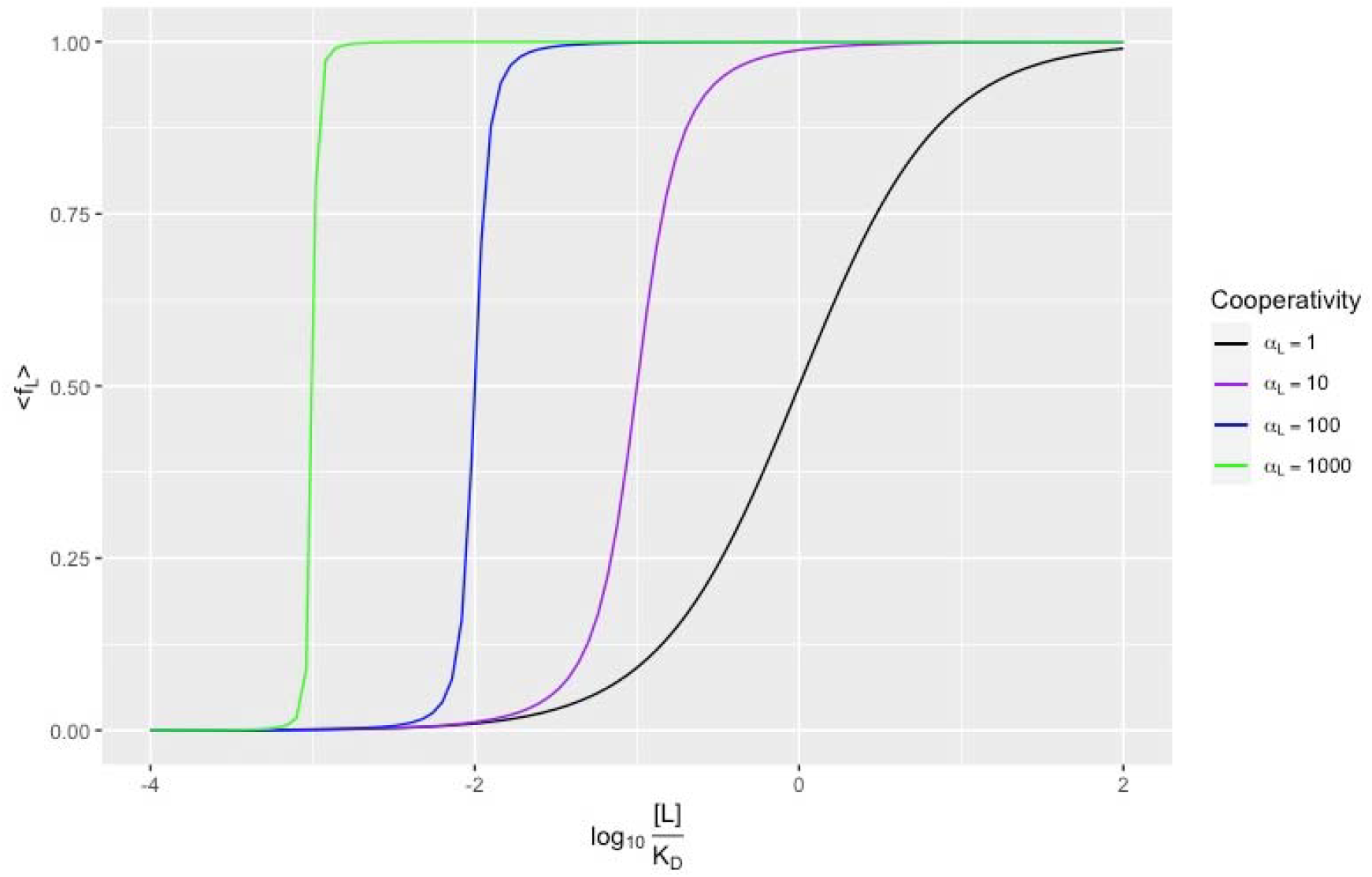
Average fraction of sites bound with radioligand for a saturation assay, assuming infinite sites, given by EQ 5. Increasing the cooperativity factor *α*_*L*_ decreases the *EC*_50,*L*_ relative to the and makes the curve steeper.

In the case of no cooperativity (*α*_*L*_ = 1), EQ 5 reduces to the familiar EQ 3. As the cooperativity *α*_*L*_ increases, the midpoint of the binding curve shifts to the left, and the curve becomes steeper. The exact expression for the midpoint is

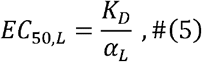

reflecting how we can increase a ligand’s binding both by improving its intrinsic binding to the protein (decreasing *K*_*D*_) and by improving the ligand-ligand cooperativity (increasing *α*_*L*_). Knowing solely the midpoint of an experimental binding curve does not allow fitting the two molecular parameters *K*_*D*_ and *α*_*L*_; we must also incorporate the shape or steepness. Comparing to the empirical Hill Equation for cooperativity, a cooperativity factor of *α*_*L*_, = 10, which has a cooperativity energy of *C*_*L*_ = −1.4 kcal / mol at room temperature, resembles a Hill Coefficient between 2 and 3 (**Fig. S3**).[25]

While we expect to see steep binding curves based on the extensive ligand-ligand cooperativity, fitting our model to published data using nonlinear regression leads to estimates of *α*_*L*_ being around 1.[29, 42] Considering most tau assays are done with *ex vivo* brain homogenate and not purified protein, the observations are typically noisy, and the confidence intervals on parameter estimates are large. Most of the published data on tau binders is on the linear (not logarithmic) scale for [*L*], making fitting to data across many orders of magnitude more difficult; this is a point to which we will return. Far more studies report radioligand displacement (competition) assays, which we consider next.

### Competition Binding Assays

#### New Parameters and Binding Isotherm

For competition experiments between a radioligand *L* and an inhibitor (ordinary test ligand) *I* at concentration [*I*], any site on the fibril can be filled by a radioligand, an inhibitor, or nothing. There are additional energies for the inhibitor’s intrinsic per-site binding Δ*G*_*I*_ (equal to 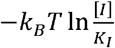 for dissociation constant *K*_*I*_), self-cooperativity *C*_*I*_, and cross-cooperativity with the radioligand *C*_*L*− *I*_ (**Fig. 4, Table 1**). Each of these cooperativity energies has an associated Boltzmann weight that multiplies the probability of a configuration: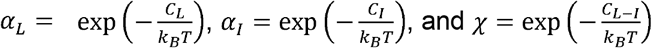.

**Table.** Symbols used in this model.

| **General Parameters for System** |  |
| --- | --- |
| $[L]$ | Free radioligand concentration |
| $K_D$ | Dissociation constant for radioligand to a single protein fibril site |
| $[I]$ | Free inhibitor (non-radioactive test ligand) concentration |
| $K_I$ | Dissociation constant for inhibitor to a single protein fibril site |
| $[F]$ | Concentration of protein fibril |
| $N$ | Total binding sites per fibril |
| $T$ | Absolute temperature |
| $k_B$ | Boltzmann's constant |
| $\lambda = \frac{\ln 2}{t_{1/2}}$ | Decay rate for radioligand with half-life $t_{1/2}$ |
| $CPM$ | Counts per minute: measured signal on detector |
| $M$ | Detection limit for voxel on detector |
| **Energetic Quantities** |  |
| $\Delta G_L = -k_B T \ln \frac{[L]}{K_D}$ | Radioligand's intrinsic binding to each fibril site |
| $\Delta G_I = -k_B T \ln \frac{[I]}{K_I}$ | Inhibitor's intrinsic binding to each fibril site |
| $C_L$ | Radioligand-radioligand self-cooperativity |
| $C_I$ | Inhibitor-inhibitor self-cooperativity |
| $C_{L-I}$ | Radioligand-inhibitor cross-cooperativity |
| **Boltzmann Factors** |  |
| $\alpha_L = \exp\left(-\frac{C_L}{k_B T}\right)$ | Radioligand-radioligand self-cooperativity |
| $\alpha_I = \exp\left(-\frac{C_I}{k_B T}\right)$ | Inhibitor-inhibitor self-cooperativity |
| $\chi = \exp\left(-\frac{C_{L-I}}{k_B T}\right)$ | Radioligand-inhibitor cross-cooperativity |
| $\beta = \frac{\chi^2}{\alpha_L \alpha_I}$ | Relative cross-cooperativity |
| **Derived Quantities** |  |
| $n_L$ | Number of sites bound with radioligand per fibril |
| $f_L = \frac{n_L}{N}$ | Fraction of sites bound with radioligand per fibril |
| $\langle X \rangle$ | Thermodynamic average of variable $X$ |
| $B_{max}$ | Maximum signal in radioligand binding assay |
| $EC_{50}$ | Concentration of ligand (radioligand or inhibitor) leading to binding of 50% the sites in a saturation assay |
| $IC_{50}$ | Concentration of inhibitor in a competition assay leading to a decrease in radioligand's signal by 50% |
| $n_c$ | Critical number of sites bound for detection |
| $n$ | Hill Coefficient |
| $k$ | Generic constant of proportionality |

**Figure 4.**
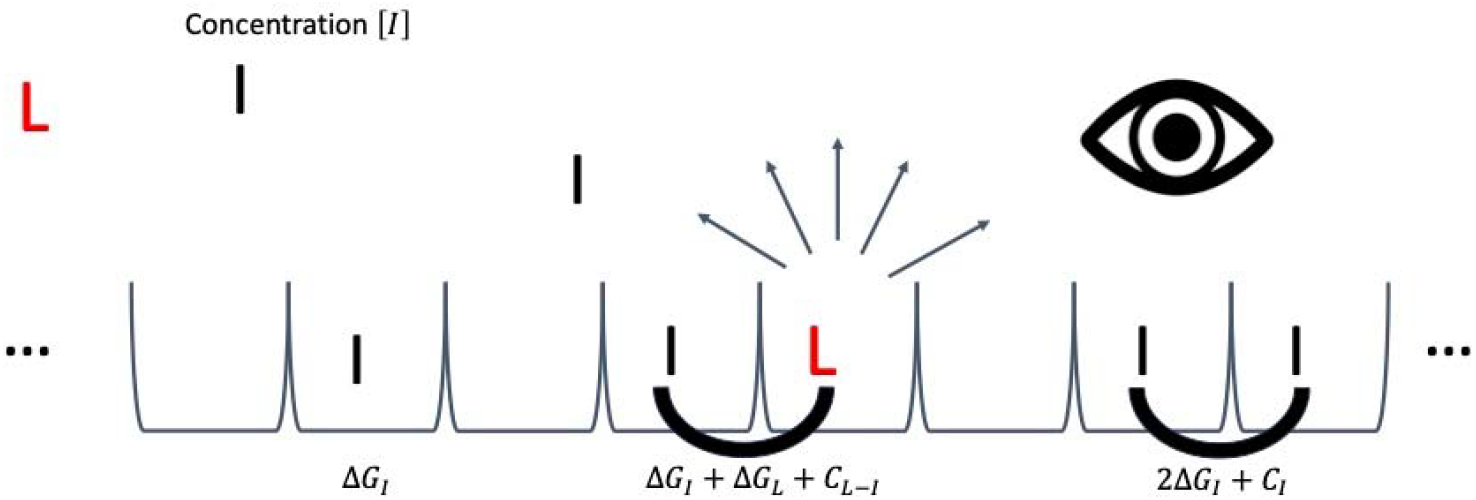
Description of a radioligand competition experiment. In addition to the parameters in a saturation assay, there is dissolved inhibitor (non-radioactive test ligand) at free concentration. The inhibitor has a dissociation constant *K*_*I*_ to each site on the fibril, changing the system’s energy by 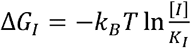 for each site bound with inhibitor. There are still nearest-neighbor interactions, adding to the system energy *C*_*L*_ for radioligand-radioligand self-cooperativity, *C*_*I*_ for inhibitor-inhibitor self-cooperativity, and *C*_*L*−*I*_ for radioligand-inhibitor cross-cooperativity. The detector directly measures binding (through radioactive decay) only for the radioligand, not the inhibitor.

**Figure 5.**
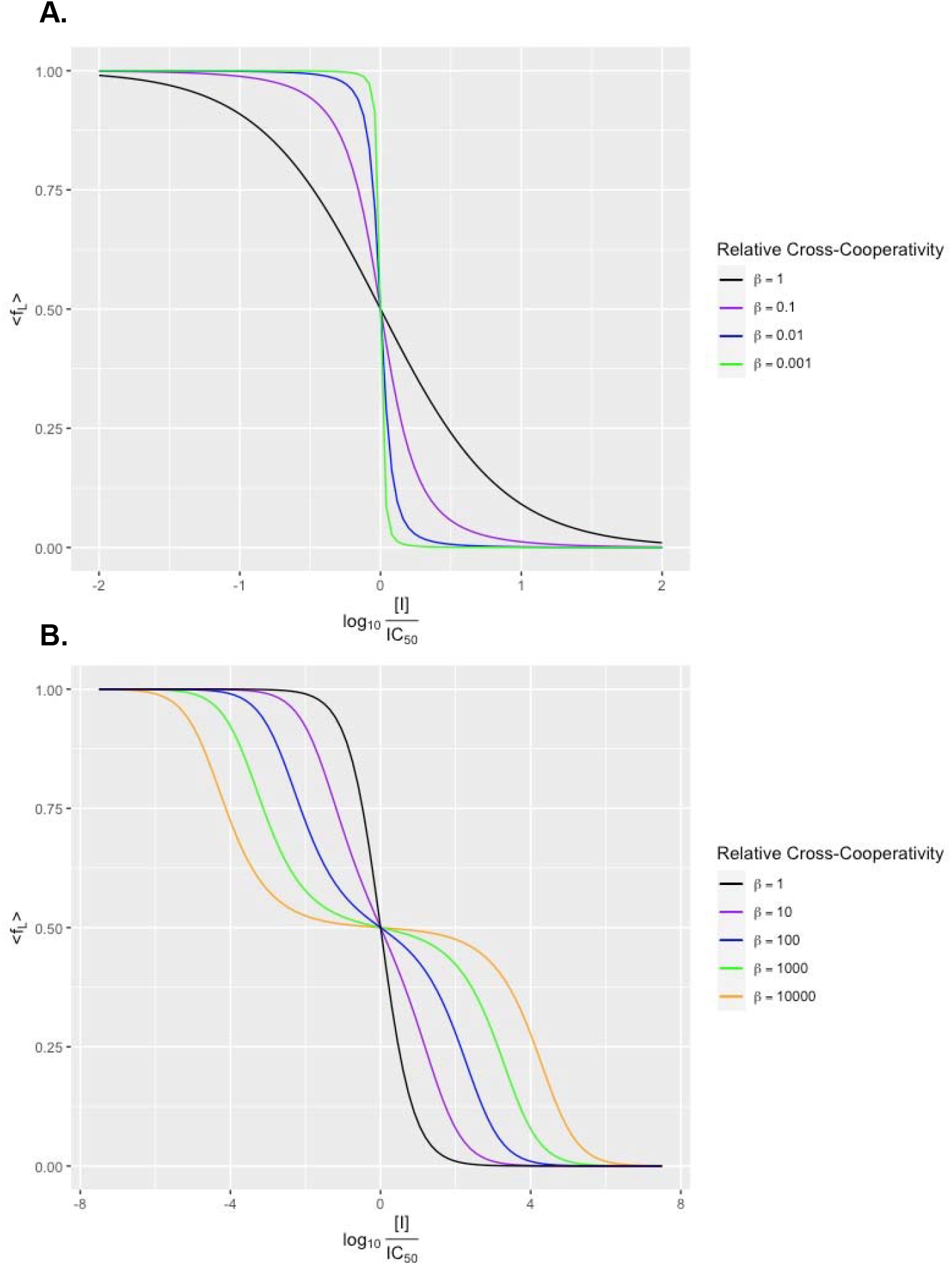
Average fraction of sites bound with radioligand for a competition assay. We assume the fibril has infinite sites that are all filled with radioligand or inhibitor and arrive at the binding isotherm given by EQ 9. The relative cross-cooperativity 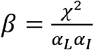 acts a shape parameter for the curve’s steepness. **a**. Plots for *β* ≤ 1. As the relative cross- cooperativity decreases, the curve becomes steeper and looks like familiar cooperative binding. **b**. Plots for *β* ≥ 1. As the relative cross-cooperativity increases, the curve becomes shallower, eventually resembling a 2-site model with a plateau about 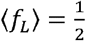 for *β* >3.

The model is now a 3-state system: each site can have a radioligand *L*, inhibitor(test ligand) *I*, or neither. Using the same transfer matrix approach with all possible parameters 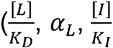, and *χ*) leads to an intractable problem of trying to find the largest of three arbitrary eigenvalues. One simplification is to assume that all the sites are bound with either the radioligand or inhibitor, reducing the problem to a 2-state system in a way that the experimentalist can control by using high concentrations of *L* and *I* (another simplification occurs when there is one common cooperativity*α*_*L*_, = *α*_*I*_ = *χ* = *α* as discussed in **Supplemental Methods**). When all the sites are bound, the fraction of sites occupied by a radioligand during a competition assay is

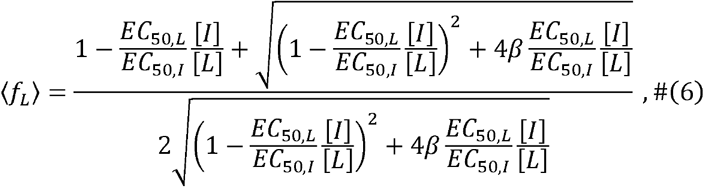

where 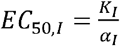 is the EC_50_ for the inhibitor binding to the fibril on its own and 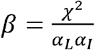 is the relative cross-cooperativity. The resemblance to the binding isotherm in the saturation assay is not a coincidence, as both come from 2-state systems (radioligand vs. empty site or radioligand vs. inhibitor). Nor is it a coincidence that the bound fraction depends only on the ratio 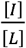: the system is saturated with radioligand or inhibitor, leaving the mole fraction of inhibitor relative to the sum of the total sites as the pertinent variable.

The midpoint of this curve occurs when [*I*] = *IC*_50_, where

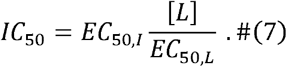

Notice the similarity to the Cheng-Prusoff Equation

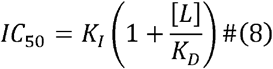

relevant for radioligand displacement assays of receptors and competitive inhibitors of enzymes following Michaelis-Menten kinetics (substituting substrate concentration [*S*] for [*L*] and Michaelis constant *K*M for *K*_*D*_).[43] For both, the *IC*_50_ increases with the radioligand concentration. Also, as the affinity of the inhibitor (*EC*_50_or *K*_*I*_) improves, the so drops, and as the affinity of the radioligand *(EC*_50_or *K*_*D*_) improves, the *IC*_50_ rises. Replacing *K*_*I*_ with *EC*_50,*I*_ and *K*_*D*_ with *EC*_50,*L*_ reflects how we can improve binding by improving intrinsic affinity to a single site and by improving cooperativity between sites. Finally, the relationship between Eqs. 7 and 8 becomes unmistakable when we realize that the +1 in Eq. 8 is not present in Eq. 7 because we assumed that the system was saturated, therefore excluding the potential for empty sites.

As a function of inhibitor concentration, the fraction of radioligand bound is then

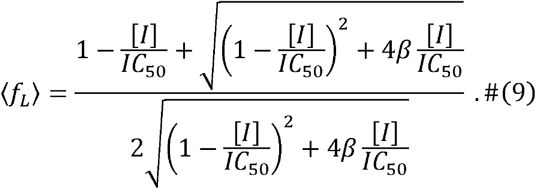

We can see that the relative cross-cooperativity *β* acts as a shape parameter when we plot the isotherm on the logarithmic scale in **Fig. 3**.

#### Non-Steep Binding Curve

This condition occurs when the relative cross- cooperativity *β* = 1, or that on average, the cooperativity exerted by the different molecular species are equal. (More precisely, the cross-cooperativity energy is the arithmetic mean of the two self-cooperativity energies: 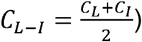. Then the identity of the neighbors does not depend on the binding energy, so each site is essentially independent. The binding isotherm should look non-cooperative, and indeed EQ 9 reduces to

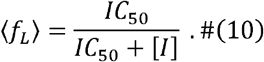

A binding isotherm like this has a steepness reminiscent of non-cooperative binding like EQ 3, yet there is clearly cooperativity in how *IC*_50_ depends on the inhibitor’s self-cooperativity *α*_*I*_ (through the dependence on *EC*_50,*I*_). An inhibitor with significant self-cooperativity, something seemingly supported by the stacked ligands of the cryoEM fibril structures, would appear to have a Hill Coefficient of *n* = 1, which we also see in binding curves.[9, 13, 14, 18, 19, 22–24] In experiments measuring displacement of a”hot” tracer by an otherwise chemically identical “cold,” unlabeled tracer, the binding parameters are identical for the ligand and inhibitor (*K*_*D*_ = *K*_*I*_, α_*L*_ = α_*L*_ = *χ* and *β* = 1 trivially). EQ 10 further reduces to the fraction of sites bound with “hot” ligand being just the “hot” ligand’s mole fraction:

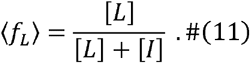

The displacement assay is uninformative for determining *K*_*D*_, giving a non-steep competition curve with the *IC*_50_ of the “cold” ligand being just the concentration of free radioligand, as we see in experimental binding curves.[14] But the procedure is still useful in validating the assumption of a single binding site in the system.

The *β* =1 condition for a competition assay reflects a general principle: in a cooperative system saturated with endogenous ligand, introducing an inhibitor leads to a non-steep competition curve. The Haldane Effect in hemoglobin is an illustrating example. Hemoglobin cooperatively binds both molecular oxygen and carbon monoxide, and since each gas causes the prosthetic heme group to absorb light at a different wavelength, binding to each can be independently measured.[44–48] In systems with carbon monoxide present, binding of oxygen is stronger: the greater the partial pressure of CO, the lower the *K*_*D*_ of oxygen.[49–52] The effect is like our model of PET tracers binding to tau in that the binding of one molecular species (like CO in hemoglobin) templates the system for the other species (like O_2_) to bind better. Unlike with the direct ligand-ligand contact in tau, the Haldane Effect is through allostery, as more of the globin subunits convert from the low-affinity tense conformation to the high-affinity relaxed conformation upon binding.[53–57] However, in both systems, as more of the system is templated with one species (radioligand for tau or CO for hemoglobin), the apparent cooperativity for the other species (test molecule or O_2_, respectively) decreases (for tau, provided that *β* = 1). Indeed, as the partial pressure of CO increases, the fitted Hill Coefficient of O_2_ decreases towards one.

#### Steep Binding Curve

When the self-cooperativities are stronger than the cross- cooperativity, then *β* <1, and the transition from the sites being mostly filled with radioligand to mostly filled with inhibitor becomes sharper. The more favorable the self- cooperativities, the smaller the value for *β*, and the steeper the curve. Because of the entropy coming from the number of binding sites, there will always be some sites with the minority molecule bound (for finite energy differences, or equivalently, for nonzero temperature), and the function never becomes a perfect step function. While we could not find examples of steep binding curves, we can imagine scenarios where designed ligands would display this behavior. A ligand-inhibitor pair could have better self- cooperativities when the plane of a flat, aromatic inhibitor binds at a different angle to the major axis of the fibril than the radioligand, so that the ligand and inhibitor sitting next to each other in the fibril is energetically frustrated. Another example would be if the ligand and inhibitor each have amide groups and can make hydrogen bonds to neighbors in the stack, but the location of the moiety on each species is different, so the geometry only works for hydrogen bonding to like molecules.

#### Biphasic (Split) Binding Curve

In many examples of radioligand displacement assays, the binding curves appear biphasic or split, which might be a result of cooperative binding to a single site on the fibril. Unlike saturation assays, in a competition experiment we can reasonably have cross-cooperativity between the two types of binding (radioligand/inhibitor or radioligand/empty) be better than the self-cooperativity, leading to *β* >1. For example, the radioligand and inhibitor can have opposite overall charges or more complementary quadrupole moments and van der Waals surfaces to each other than to themselves. In this case, substantial inhibitor binding occurs further below the *IC*_50_ than in the non-cooperative model. An apparent transition occurs for *β* >3 (**Fig. S4**): there is a plateau around 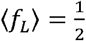 that becomes flatter and wider as *β* becomes larger. We can justify this by observing that going down the length of the fibril, most of the sites are alternating between the radioligand and inhibitor being bound (**Fig. S5**).

#### Fitting the Model to Data

An experimenter can fit this model to data using nonlinear regression, for example with the “nls” function in the R Programming Language.[29, 42] The full set of molecular parameters is over-determined given the available data in a competition assay. Unless one knows *K*_*D*_ and *α*_*L*_ from a previous saturation assay of a given radioligand, one cannot decompose *IC*_50_ into *EC*_50,*L*_ and *EC*_50,*I*_ (the experimenter can always control [*L*]). They also cannot break apart *β* into the different cooperativities. Even if the experimenter does know the molecular parameters for the radioligand, they would not be able to separate *EC*_50,*I*_ into *K*_*I*_ and *α*_*I*_, or *β* into *α*_*I*_ and *χ* (knowing *α*_*L*_). The assay is still practical: the measured *EC*_50,*I*_ tells us the inhibitor’s affinity. We just do not know how much of the effect comes from per-site binding and how much comes from cooperativity. Even knowing just the *IC*_50_ values for different inhibitors in identical assay setups can tell us which one has a better *EC*_50_. EQ 7, applied to inhibitors 1 and 2, tells us

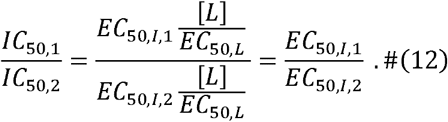

### Detection Limit in PET Experiments

The entropy from the number of binding sites and cooperativity between sites affect a fibril’s ability to appear in PET scans. Assume that a detector can only measure positron emission for a region of space if the counts per unit time coming from that space is at least *M*. Assume the radioligand has decay rate *λ*, the fibril has *n*_*L*_ sites bound with radioligand, and the tissue has concentration of fibril [*L*]. Then detection occurs if *λn*_*L*_[*F*] ≥ *M*, or if

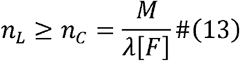

for critical value *n*_*c*_. This condition applies for detection at a single time point; if the measurement happens at a longer time scale than thermodynamic equilibration of radioligand binding, then the condition is ⟨*n*_*L*_⟩ ≥ *n*_*c*_. Then the average number of sites bound increases with the fraction of sites bound (*f*_*L*_, dependent on intrinsic binding affinity *K*_*D*_ and cooperativity *α*_*L*_) and the total number of sites *N*.

For example, if every fibril has the same number of sites *N* and the radioligand has no cooperativity (*α*_*L*_=1), then the number of bound sites per fibril follows a binomial distribution (*N* independent trials, each with probability of success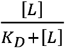). [58] The average number of sites bound would be 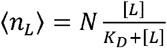, and the condition for detection would be

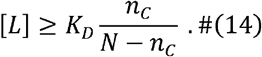

Since *N* is always larger than *n*_*c*_, we see that the necessary free ligand concentration for a fibril to appear on a PET scan can be much lower than the *K*_*D*_, even in the absence of cooperativity. In other cases of drug development, we need the *in vivo* concentration of the drug to be higher than the *K*_*D*_ so that enough of the enzyme gets inhibited or enough of the receptor gets activated, for example. A PET scan is fundamentally a labeling problem, and since each tau fibril has many sites for possible labeling, a useful tracer need not have the affinity expected of other drugs (even if the cooperativity between sites is helping substantially).

## Discussion

Motivated by the appearance of atomic resolution structures of ligand/fibril complexes, we propose a new model of ligand binding that considers entropy from the number of sites and cooperativity between sites. Using only a nearest-neighbors (Ising) mode of cooperativity, this model better accounts for the available binding data than a non-cooperative model. Despite having measured affinities at or below the nanomolar range, the ligands make far more contacts with each other than with the protein, and the ligand-ligand contacts seem to be ideal for π-π stacking through permanent quadrupole moments. These features of ligand binding suggest that ligand-ligand cooperativity plays a large role in determining experimental affinities, and EQ 5 reflects this, describing *EC*_50,*L*_ in terms of intrinsic per-site binding and cooperativity. Experimental neighbors radioligand competition curves tend to have the steepness associated with a Hill Coefficient of *n*=1 or appear to have 2 sites for binding. Nearest-cooperativity in a system saturated with radioligand and inhibitor can explain both these curve shapes by having different radioligand-inhibitor cross-cooperativities.

This model nonetheless has limitations. Even though our model for a competition experiment allows for steep or biphasic curves, we so far have only found examples that do not look steep. The non-steep curves may be expected, however, since most of the known tau binders are similar (large, flat, hydrophobic, fused aromatic heterocycles), making the self-cooperativities and cross-cooperativities roughly equal (and *β* ≈ 1). One could argue that the unusual binding curves and affinities result from multiple inequivalent binding sites along a single fibril. Multiple modeling studies have suggested secondary binding sites outside of each protofilament’s main trough (or the inter- protofilament cleft for the disaggregator EGCG).[59, 60] The structural data, meanwhile, only has diffuse, unassigned density at other sites. The existing computational fits to multiple binding sites do not model multiple ligands, let alone cooperativity between ligands. Similarly, this work does not rule out more complicated models of cooperativity (e.g., delocalized coupling along the fibril axis), but this simpler model is sufficient to explain available data.

Notwithstanding these caveats, this model of cooperativity and entropy is consistent with available structures, affinity data, and practical effects for PET scans. It also generalizes to other cooperative systems.

## ASSOCIATED CONTENT

## Supporting information

Supporting Information is available free of charge on the PNAS Publication website: Figures S1–S5 and Supplemental Methods

## Author information

### Authors

**William F. DeGrado** - Department of Pharmaceutical Chemistry and Cardiovascular Research Institute, University of California, San Francisco, 94158, United States.

**Michael Grabe** - Department of Pharmaceutical Chemistry and Cardiovascular Research Institute, University of California, San Francisco, 94158, United States.

**Brian K. Shoichet** - Department of Pharmaceutical Chemistry, University of California, San Francisco, 94158, United States.

## Author Contributions

Conceived by MSS and BKS. Development of the model, calculations, and MCMC simulations by MSS. Model refined by MSS, WFD, MG, and BKS. All authors contributed to writing and editing the manuscript.

## Notes

The authors declare the following competing financial interest(s): BKS is co-founder of BlueDolphin LLC, Epiodyne Inc, and Deep Apple Therapeutics, Inc., and serves on the SRB of Genentech, the SAB of Schrodinger LLC, and the SAB of Vilya Therapeutics. WFD is a member of the scientific advisory boards of Alzheon Inc., Pliant, Longevity, CyteGen, Amai, and ADRx Inc., none of which contributed support for this study.

## Abbreviation Used

AD: Alzheimer’s Disease
CBD: corticobasal degeneration
*CPM*: counts per minute
Cryo- EM: cryogenic electron microscopy
CTE: chronic traumatic encephalopathy
*EC*_50_: effective constant of 50%
EGCG: epigallocatechin gallate
*I*: inhibitor
*IC*_50_: inhibitory constant of 50%
*L*: (radio)ligand
*LR*: (radio)ligand-receptor complex
NFT: neurofibrillary tangle
PET: positron emission tomography
PHF: paired helical filament
PSP: progressive supranuclear palsy
*R*: receptor
SASA: solvent-accessible surface area

## Supplemental Information

**Figure S1.**
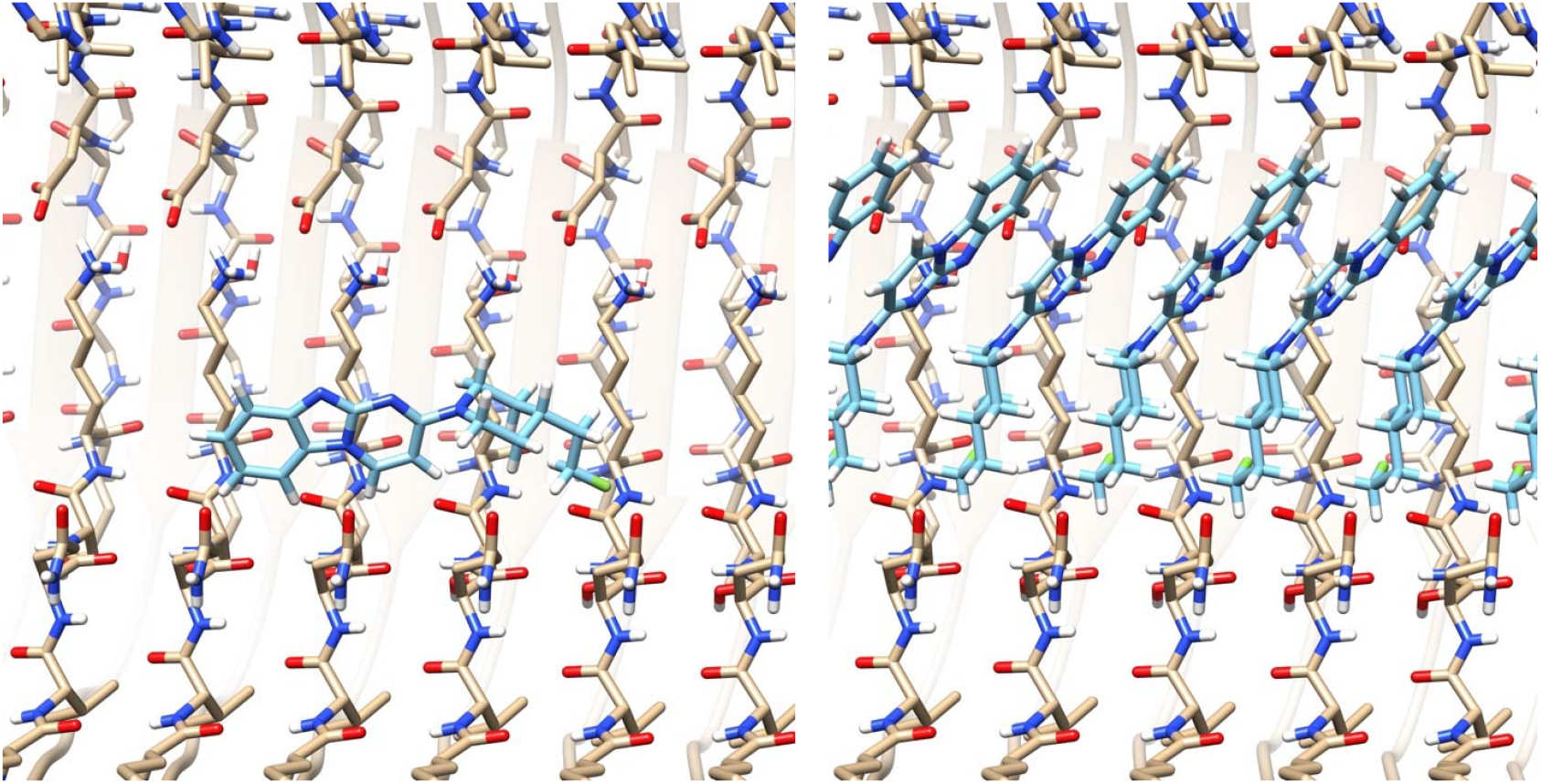
Docking the molecule GTP-1 with (left) and without (right) a stacking symmetry requirement shows how the molecule’s interactions with the AD PHF tau fibril change.

**Figure S2.**
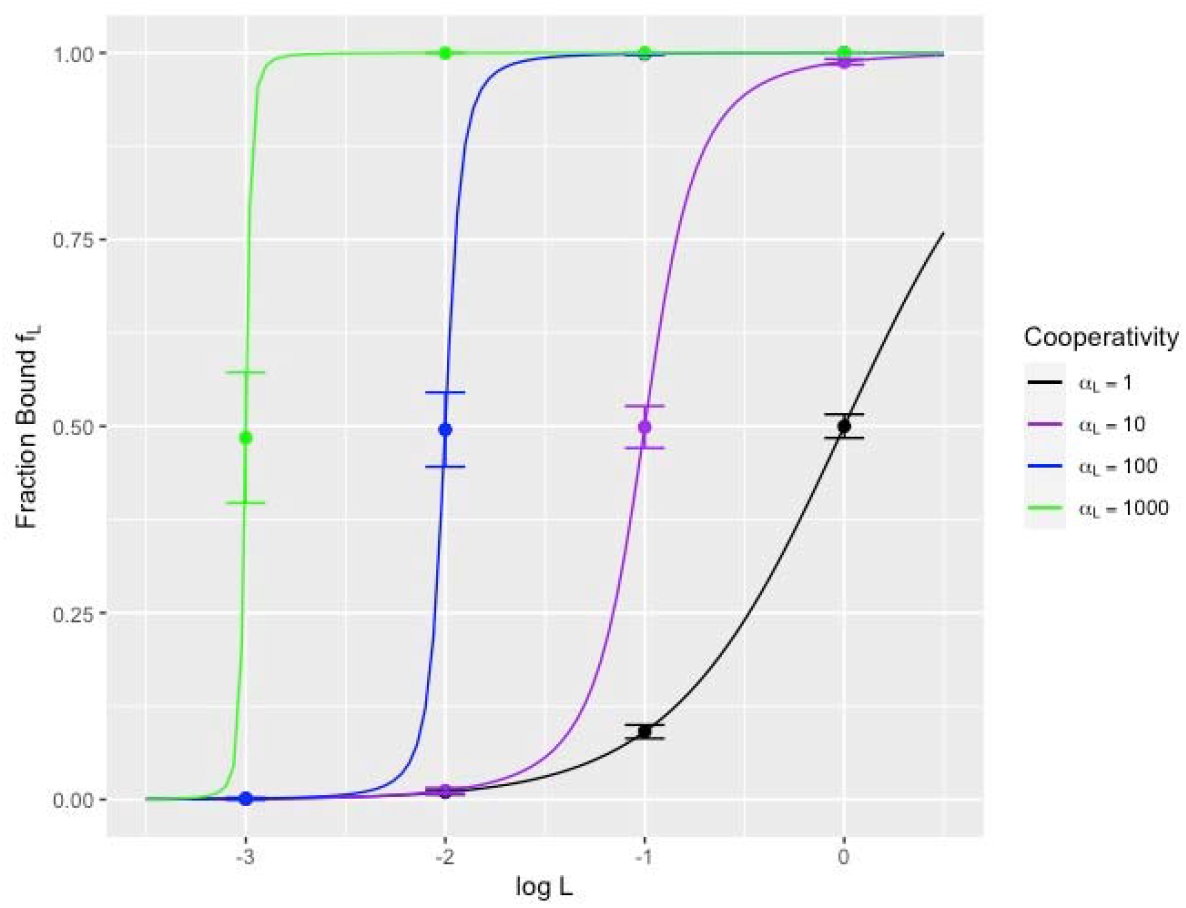
Theoretical binding curves from a saturation experiment for binding to an infinite symmetric fibril compared to MCMC simulations run for a fibril of length *N* = 1000 at different (*L,α*_*L*_) values. The MCMC simulations comprised modeling the fibril of a1000-bit binary string, where each proposed step was flipping a random bit with acceptance probability given by the Boltzmann factors. Each (*L,α*_*L*_) pair had 10 independent MCMC simulations run at different seeds for the initial configuration and pseudo-random number generator. We ran 1 million steps of equilibration after initializing each simulation without collecting data to equilibrate. After equilibration, we collected data (the binary string) for 10 billion simulation steps. Points on the graph denote averages and error bars denote standard deviations of the aggregate counts across all 10 simulations seeds. Using 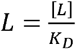.

**Figure S3.**
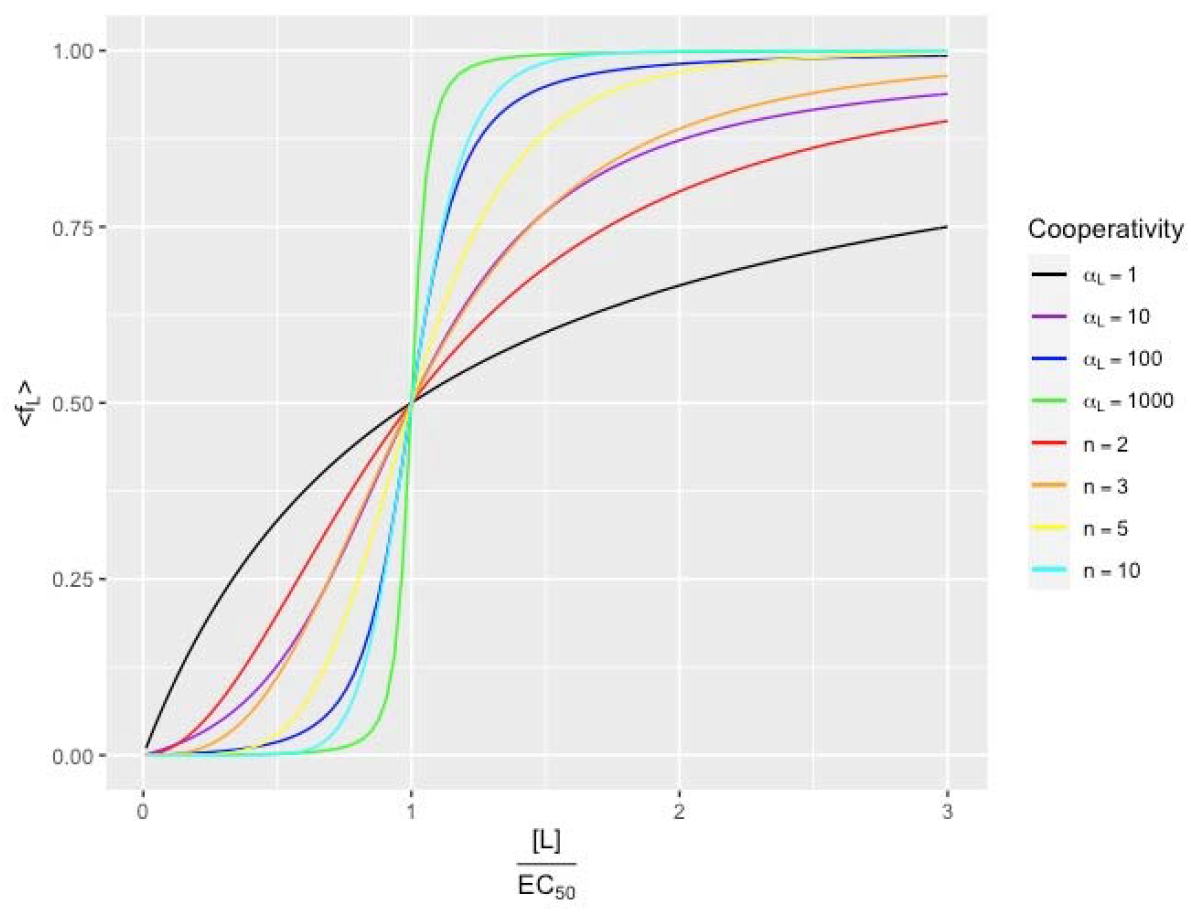
Comparing the cooperativity between the infinite symmetric fibril model in a saturation experiment (for different cooperativity factors *α*_*L*_) to the empirical Hill Equation

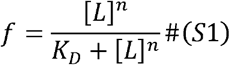

(for different Hill Coefficients *n*). For the infinite symmetric fibril, 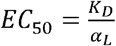 by EQ 5 in the main text, and *K*_*D*_ has units of mol / L. The Hill Equation gives the fraction bound with no ligand depletion for the reaction *R* + *n*_*L*_ ⇌ *RL*_*n*_ (for receptor *R* and ligand *L*), so *K*_*D*_ has units of (mol / *L*)^*n*^, and 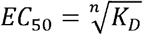.

**Figure S4.**
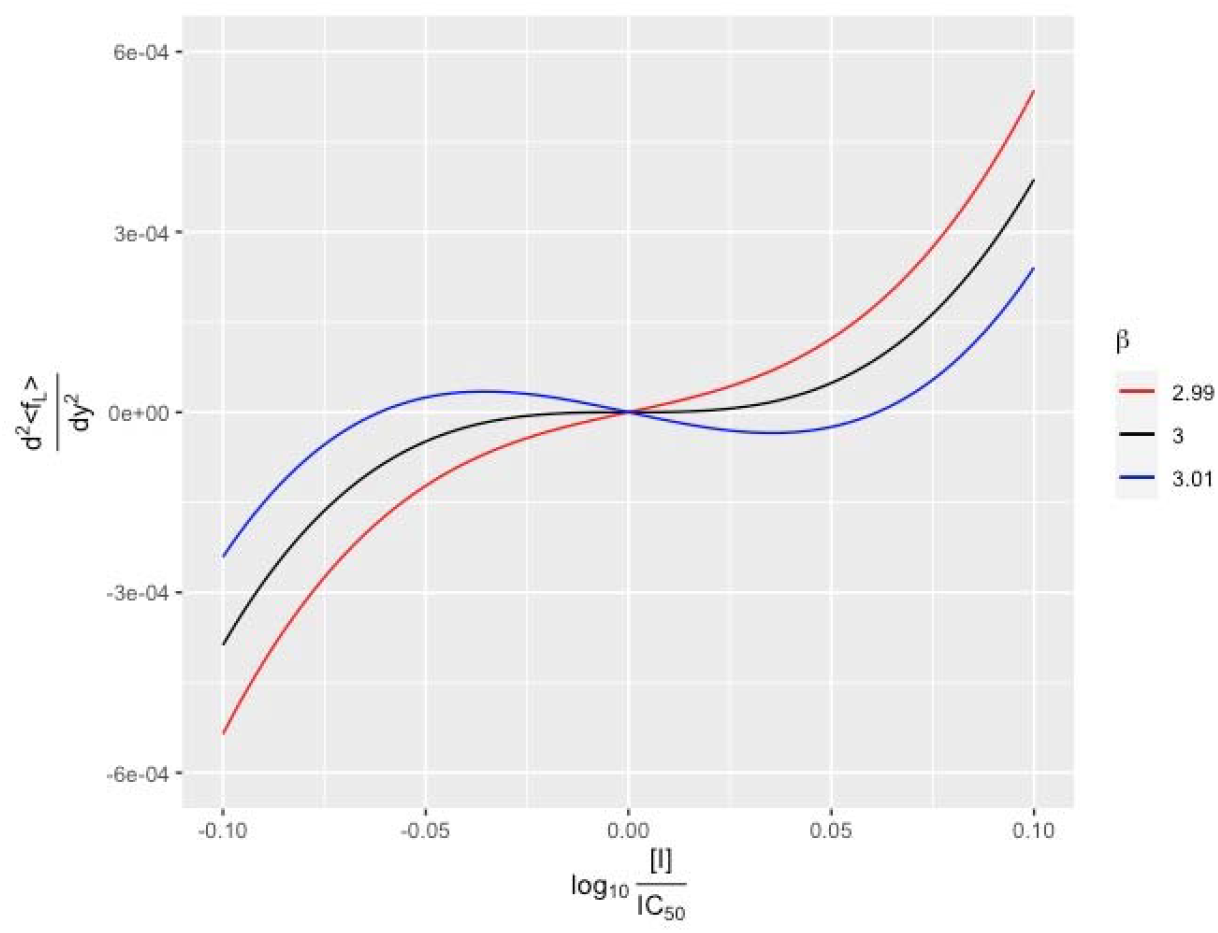
Showing the transition from monosphasic to biphasic competition binding curves at *β* = 3 by plotting EQ S36.

**Figure S5.**
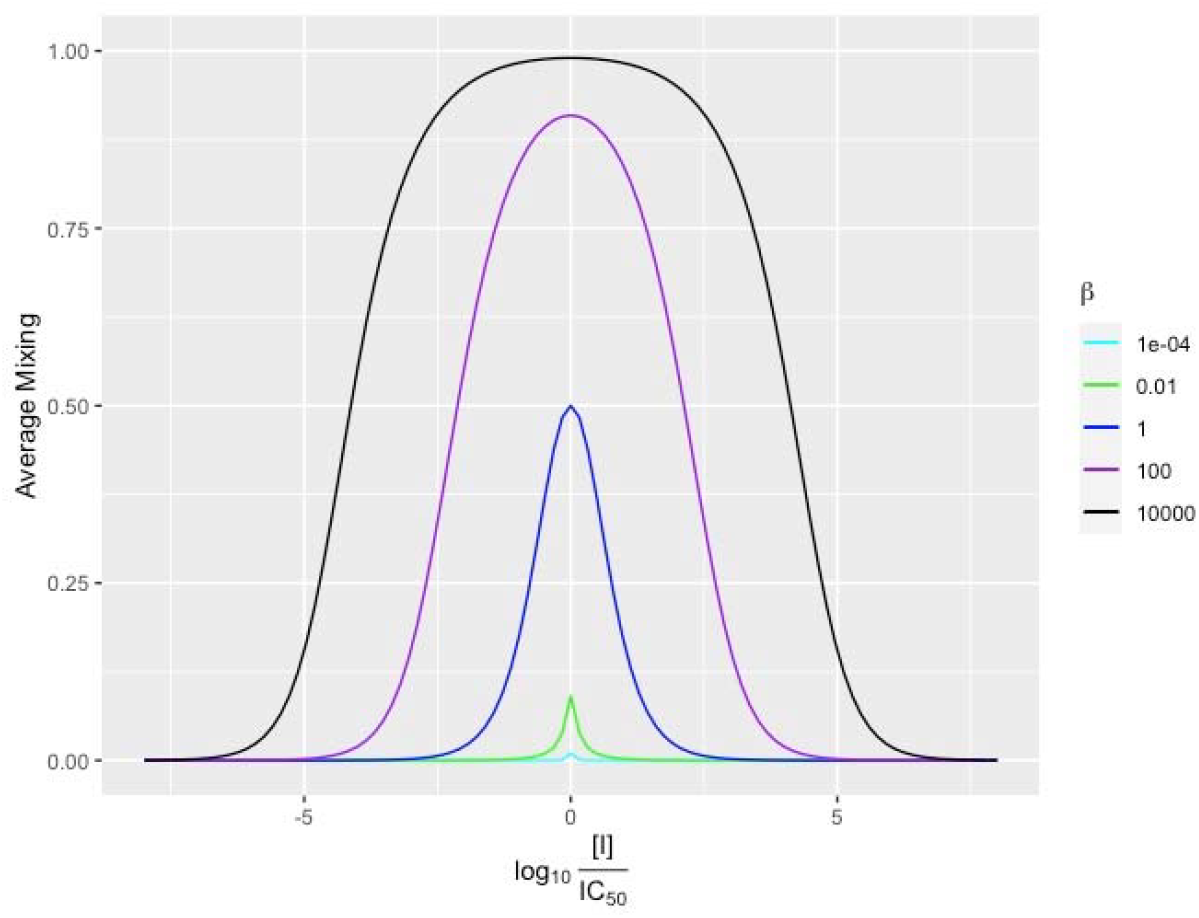
Average mixing between sites bound with radioligand and sites bound with inhibitor in the competition assay when all sites are bound with something as given by EQ S43.

### Supplemental Methods: Transfer Matrix for Equilibrium Isotherm

We will begin with deriving the equilibrium isotherm for the radioligand saturation has experiment. A fibril with *N* sites, each of which can be filled with a radioligand or empty, has 2*N* possible configurations. Each configuration has an equilibrium probability proportional to its Boltzmann weight:

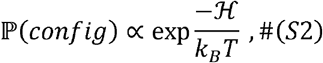

where *K*_*B*_ is Boltzmann’s constant, *T* is the absolute temperature, and ℋ is the energy (Hamiltonian) of the configuration. The system gets an added energy 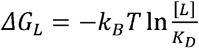 for each site occupied with radioligand, for free ligand concentration [*L*] with dissociation constant *K*_*D*_, assuming no ligand depletion. There is additional energy coming from Introduce nearest-neighbors (Ising) cooperativity *C*_*L*_. To enumerate the sites on the fibril, and therefore all the configurations, label the sites in the linear array from 1 to *N*. two empty sites on either end of the array labeled 0 and *N* + 1; this will make the enumeration easier without affecting the total number or energy of each configuration. Then with indicator function *s*_*i*_ taking value 1 if site *i* is occupied and value 0 if site *i*is empty, the energy of a configuration is

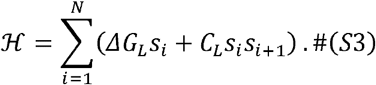

The partition function is the constant of proportionality for the probabilities

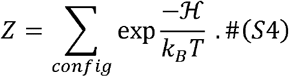

Let *n*_*config*_ be the number of sites bound for a given configuration. Then the thermodynamic average of the number of sites bound ⟨*n*⟩ can be written as

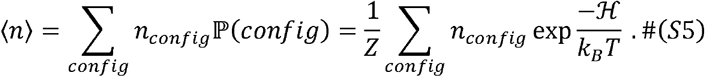

Notice that the Hamiltonian is only a function of the number of sites bound *n*_*config*_ and the number of occupied neighboring pairs of sites *p*_*config*_:

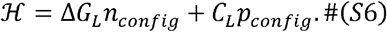

So using 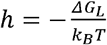 and 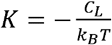 allows us to write the Boltzmann weight as exp(*hn*_*config*_, + *Kp*_*config*_) the partition function as

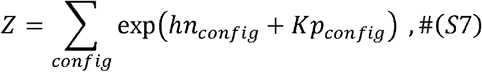

and the average number of sites bound as

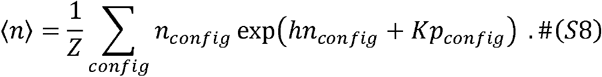

In the same way one can get the average energy by taking the appropriate partial derivative of In *Z*, notice that

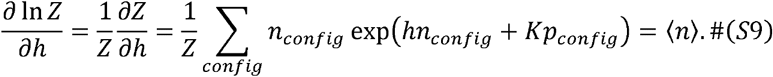

We can further simplify by using 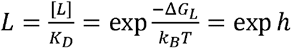 no brackets on the *L* to indicate non-dimensionalized). Using the Chain Rule from calculus, we finally arrive at

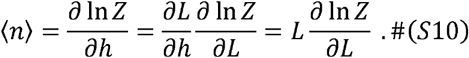

So, like much of statistical mechanics, this problem reduces to finding the partition function.

There is a technique from condensed matter physics for finding the partition function of an infinite linear array of interacting particles: the transfer matrix. In our case, each position on the fibril *i* can be in one of two states, denoted in Dirac Notation by the kets 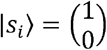 for a filled site and 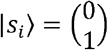 for an empty site. In this basis, the transfer matrix element ⟨*s*_*i*+1_ |*T*|*s*_*i*_ ⟩gives the extra Boltzmann weight in going down the fibril from site *s*_*i*_ to site *s*_*i*+1_. That is,

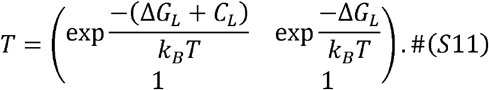

The calculations will be easier if we split the Boltzmann weight in going from an empty site to a filled site (or vice-versa) into two equal parts, making the matrix symmetric:

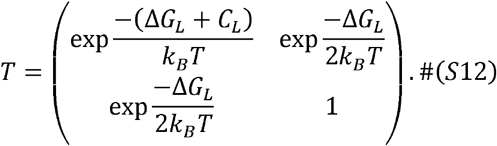

In terms of *L* and *α*_*L*_, the transfer matrix is

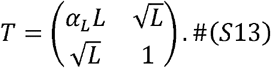

Using the fact that the extra sites *s*_0_ and *s*_*N*+1_appended to each end are empty, several resolutions of the identity, and a proof by induction, we can write the partition function as

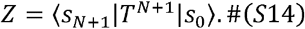

Since *T* is Hermitian by virtue of being real and symmetric, the Spectral Theorem of linear algebra guarantees that it has an orthonormal basis of eigenvectors and real eigenvalues:

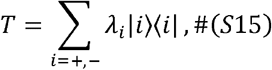

where *λ*_+_ is the larger eigenvalue with normalized eigenvector |+⟩ and *λ*- is the larger eigenvalue with normalized eigenvector |−⟩. Using the orthonormality of this basis, we can write

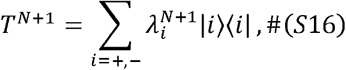

Since 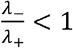, in the limit of large *N*, we have

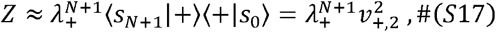

where 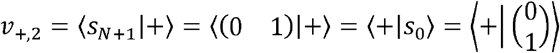 is the second component of the normalized eigenvector with eigenvalue 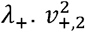 is of order 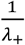, so we get the final simplifications

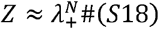

and

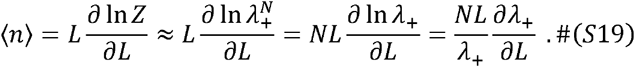

The eigenvectors of the matrix *T* are

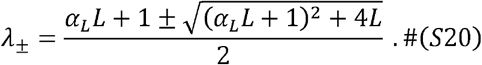

Plugging the expression for *λ*_+_ into EQ S19 and simplifying, we get

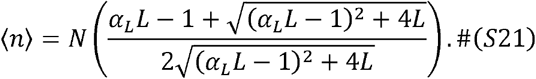

Assuming that the fibril is long enough that there are no significant edge effects on the many sites in the middle, we have ⟨*n*⟩ = ⟨*Nf*⟩ = *N*⟨*f*_*L*_⟩for the average of the fraction of sites bound with radioligand *f*_*L*_. Then

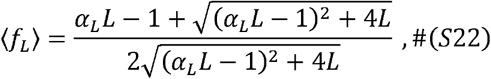

which is the binding isotherm for a saturation assay assuming no ligand depletion.

There is another way to calculate this isotherm from the partition function that will be useful for the case of competition binding assays. For an arbitrary site *i*, its fraction bound (thermodynamic average over 0 and 1 over all configurations) is

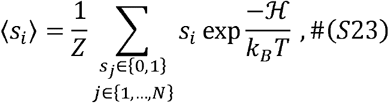

meaning that we evaluate the occupancy of site *i* (0 for empty and 1 for filled) for each configuration, weight the value by the Boltzmann weight of the configuration, sum, and normalize by the partition function.

Introduce the “pseudo-Pauli matrix”

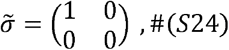

named so because it serves the same purpose as the Pauli matrix 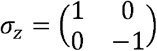 in the theory of the Ising magnet but does not have zero trace. As with *σ*_*z*_ in the Ising magnet, the action of 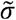 on the state ket of site *i* serves to multiply by the state occupancy:

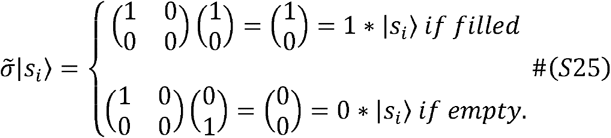

By an argument like the one leading to the expression for the partition function in EQ S14, we can write EQ S23 for the average ⟨*s*_*i*_⟩ as

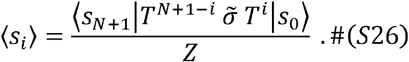

If the site of interest is in the middle of a sufficiently long fibril, then both *i* and *N* + 1− *i* will be large, so 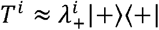 and 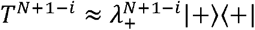. Then

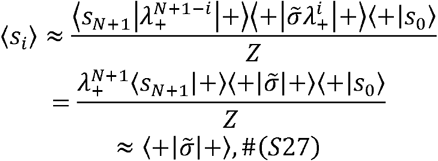

since 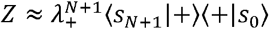 by EQ S16. Evaluating this matrix element is equivalent to squaring the first component of the normalized eigenvector of the larger eigenvalue and gives the same expression as EQ S22.

We can use the same approach to derive the equilibrium isotherm for a competition experiment. In this case, each site can have a radioligand bound, have an inhibitor (test ligand) bound, or be empty. This situation describes a 3-state system: in

Dirac notation, the kets become 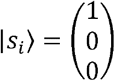 for radioligand binding, 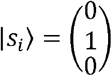 for inhibitor binding, and 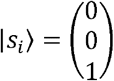 for empty. There are similar parameters involving the inhibitor as with the radioligand, as described in the main text: 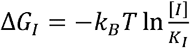 added for each site bound with inhibitor at free concentration [*I*],*C*_*I*_ = − *K*_*B*_*T* In *αI* added for every pair of neighboring sites both bound with inhibitor, and *C*_*L*−*I*_ *K*_*B*_*T* In *χ* for each pair of neighboring sites bound with different species. The transfer matrix is then

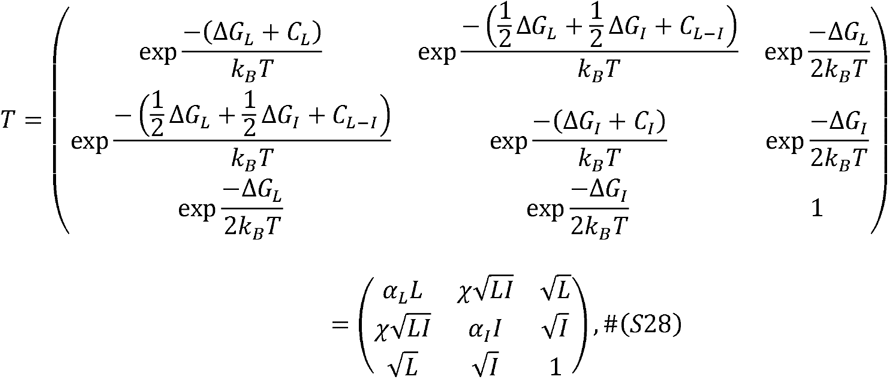

where 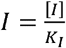. We have again broken the Boltzmann weight contribution for going from one site to one with a different occupancy (off-diagonals) in two so that the matrix will be symmetric (and therefore be diagonalizable with real eigenvalues). There will again be two appended empty sites: 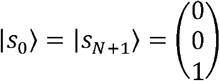 The partition function is still 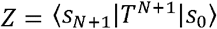. Assuming *T* is non-degenerate, call the largest of its 3 eigenvalues *λ*_1_, with normalized eigenvector |1 ⟩. We can similarly find the average fraction of sites bound with radioligand as 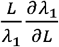 EQ S19) or 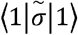(EQ S27), using a new 3 × 3 matrix 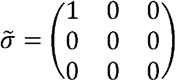.

While the saturation assay gave a 2-state system, a 2 × 2 transfer matrix, and 2 eigenvalues easily found with the quadratic formula, this setup of the competition assay requires finding the largest of 3 eigenvalues of an arbitrary real symmetric matrix. Assuming every site is bound is one way to reduce this problem to the tractable 2-state system. The transfer matrix looks like

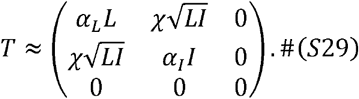

We can see that this approximation is valid at concentrations of radioligand and inhibitor high enough, given their molecular parameters, so that *α*_*L*_*L; α*_*I*_*L*, and 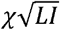 are each much larger than 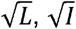 and 1. Even if we do not know the molecular parameters *K*_*I*,_ *α*_*L*_, *α*_*I*,_ and *χ a priori* (hence the experiment), we can always increase the free ligand concentrations (up to solubility limits) so that this approximation works. In this limit, 0 is obviously an eigenvalue, and the other two are 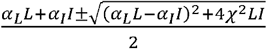, so the largest is the non-zero one with the plus sign. Then by either method, we can get the expression for ⟨ *f*_*L*_ ⟩ given by EQ 6 in the main text.

Another way to simplify the transfer matrix ins EQ S28 is to assume that there is one common cooperativity between all pairs of molecular species: *α*_*L*_ = *α*_*I*_ = *χ = α* means

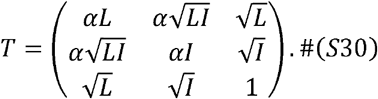

While the experimenter does not have control over the system to ensure this condition is always met, an approximation along the lines of *α*_*L*_ ≈ *α*_*I*_ *χ* is reasonable if both species are very similar. The approximation is always exact if the two species are “hot” (radiolabeled) and “cold” versions of the same molecule, which the experimenter can control. Unlike in the condition before, we do not need to have every site be filled, meaning we can run the experiment with arbitrary radioligand and inhibitor concentrations. The transfer matrix in EQ S30 is just simple enough that we can find its largest eigenvalue: 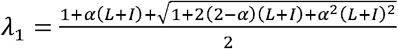. Using the approach of 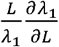(EQ S19) or 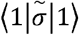(EQ S27), we then have

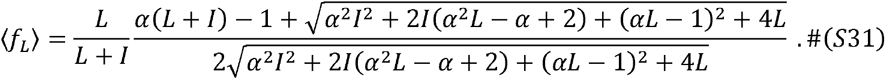

This function has all the expected limits. First, filling all the sites by making *α* ≫ 1 (equivalently, making the transfer matrix in the derivation into 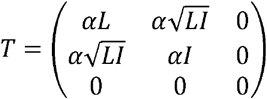 leads to

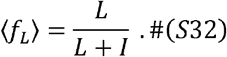

We also get this limit by first imposing the requirement that every site is filled, then deriving EQ 9 in the main text, and then imposing *α*_*L*_ = *α*_*I*_ *χ =α*. The meaning of this limit is that each site can be bound with radioligand (Boltzmann weight *α*^2^*L*) or inhibitor (*α*^2^*L*), and the constant cooperativity makes the binding across sites independent, giving 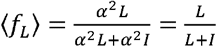. We also get the halfway point when *I* = *L*, or

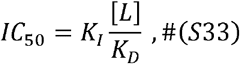

which agrees with the main text’s EQ 7 when *α*_*L*_ =*α*_*I*._

The next limit to check is *α*_*I*_ = 1, meaning there is no cooperativity between any pair of molecular species. Then

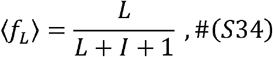

which is what we would expect for competitive inhibition of independent receptors or enzymes operating with Michaelis-Menten kinetics. The halfway point is exactly the Cheng-Prusoff Equation

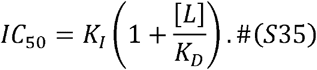

Finally, adding no inhibitor to the system means setting *I* = 0, leading to the saturation assay binding isotherm given by EQ 5.

For the case of every site being bound, we can derive the biphasic behavior for *β* > 3. We define a biphasic isotherm as one with 3 inflection points on the logarithmic axis. With 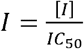 and *y* = log_10_ *I*, an inflection point occurs whenever 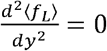 for a given *β* value. Using the expression for⟨*f*_*L*_⟩ in EQ 9 in the main text,

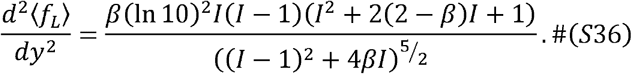

Since *I* > 0 the denominator is always positive, and the inflection points are given by the roots of the numerator. We never get the root at *I* > 0 and always get the root at *I* > 1 (the *IC*_50_). The quadratic in the numerator has roots at 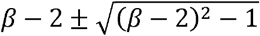. These roots are real and positive only for *β* > 3. The case of *β* = 3 just increases the multiplicity of the root at 1. If the cooperativity energies *C*_*L*_, *C*_*I*_ and *C*_*L*−1_do not depend on temperature, then the phase transition at *β* = 3 happens at temperature

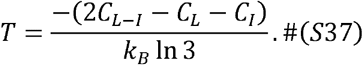

Lastly, we derive the average mixing for the competition assay as follows. First, allow for a site to have either molecular species or nothing bound. Use the indicator function *s*_*i*_ for whether a site has radioligand bound and the indicator function *i* for whether a site has inhibitor bound. Since a site cannot have both species bound at the same time, the possible states for the system described by 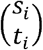 are 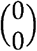 for empty,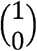 for radioligand bound, and 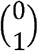 for inhibitor bound, with 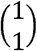 not allowed. This scheme describes the same 3-state system as before, and the extra empty sites appended to either end of the fibril means *s*_0_ = *s*_*N*+1_ = *t*_0_ = *t*_*N*+1_ = 0. The mixing across all sites of a given configuration is the number of radioligand sites followed by inhibitor sites plus the number of inhibitor sites followed by radioligand sites: 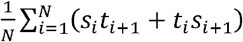. The energy (Hamiltonian) for a given configuration is

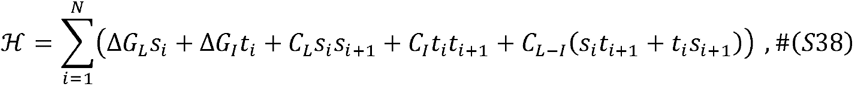

the partition function is

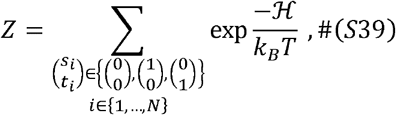

and the (thermodynamic) average mixing is

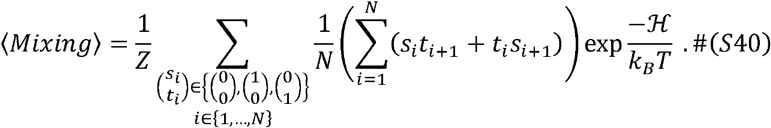

If we define 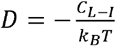, so 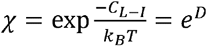, then we can find the average mixing as

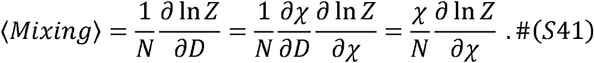

If we use the assumption that all the sites are bound with something (radioligand or inhibitor), then we get the simplification of 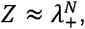 where 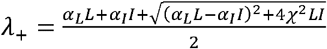.

Then

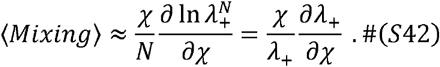

After simplifying with 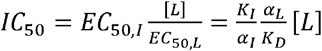 (from EQ 5 and EQ 7 in the main text) and 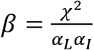,we get

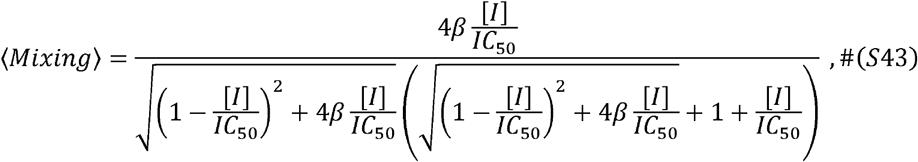

which we plot above as a function of 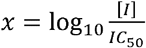 for different values of *β*. The mixing is always the best at [*I*] = *IC*_50_, and as *β* increases, the range of inhibitor concentrations for good mixing increases. The mixing will only be perfect (equal to 1) in the limit of infinite *β*, which can happen at zero temperature, at which case the entropy of mixing does not contribute to the free energy.

